# The frequency of resistant *Escherichia coli* isolation from seagulls is positively related to human population density

**DOI:** 10.64898/2026.08.03.742510

**Authors:** Rebecca Abraham, André Becker Saidenberg, Kittitat Lugsomya, Shewli Mukerji, Marin Milotic, David Jordan, David J. Hampson, Marc Stegger, Sam Abraham

## Abstract

Antimicrobial resistance (AMR) is a major global public health threat. Wild birds, including seagulls, are increasingly recognised as potential reservoirs and disseminators of resistant bacteria linked to human activity. The objective of the study was to assess the association between human population density and the occurrence of *Escherichia coli* resistant to critically important antimicrobials in Australian seagulls. Faecal samples were collected from seagull populations in coastal regions across Australia representing differing human population densities. Resistant *E. coli* isolates were identified and characterised using multilocus sequence typing and plasmid incompatibility group analysis to determine relatedness to human associated lineages. The frequency of resistant *E. coli* isolation increased with human population density. The predominant sequence types ST10, ST131 and ST354 comprised 24.5% of isolates and belong to globally distributed human associated lineages linked to extraintestinal pathogenic *E. coli*. Many isolates carried IncF and IncI plasmids, which are key vectors of *bla*_CTX-M_ extended spectrum beta lactamase genes and plasmid mediated quinolone resistance determinants commonly reported in human clinical strains.

**IMPORTANCE:** These findings support the contention that seagulls primarily acquire resistant bacteria through contact with anthropogenic activities. Once acquired, these bacteria may be disseminated to other seagulls, birds and animals, including being transmitted to humans.

## INTRODUCTION

The development of antimicrobial resistance (AMR) by bacterial pathogens reduces options for treatment, and this has become a global one-health issue with significant ramifications for public health. Transmission of AMR amongst bacteria, and ultimately to pathogenic species and strains, may occur across a network of pathways linking humans, animals, food, and the environment. Knowledge of the extent of risk posed by specific pathways is typically sparse, in part because of the vast number of combinations of host, drug, bacterial agent and other factors that comprise each pathway, but also because many routes of transfer occur within natural ecosystems that do not easily allow for testing of hypotheses. Nevertheless, where these pathways can be studied, emphasis is required on the extent of resistance to critically important antimicrobials (CIA), including carbapenems, 3^rd^ generation cephalosporins, polymyxins, and quinolones, since these form the last line of defence in clinical settings ^1^, and the early detection of such resistance is paramount ^2^. Similarly, where specific vertebrate hosts occupy an ecological niche that on a priority basis could exacerbate the propagation of resistance to CIAs (CIA-R), these should be a focus for attention. In both these respects, wild birds are of special interest since their mobility across large geographic areas enhances the potential for transmission from and to other hosts (perhaps indirectly through the environment), and in some instances their ecology brings them into intimate contact with human populations. Moreover, there is substantial evidence in the international literature of wild birds being colonised by bacteria exhibiting CIA-R ^3–5^. Several studies have shown that various species of seagulls readily acquire and disseminate resistant bacteria into the wider environment due to their interaction with aquatic and terrestrial environments, and willingness to expose themselves to anthropogenic activities ^6–9^. In an Australia-wide survey conducted between 2016 – 2017, a high prevalence of faecal carriage of CIA-R was uncovered in *E. coli* from silver gulls (*Chroicocephalus novaehollandiae*), particularly involving fluoroquinolone and extended-spectrum cephalosporin-resistant *E. coli*, with reported prevalences of shedding of 24% and 22% respectively ^10^. Another study in Australia examined 425 *E. coli* isolates from silver gulls and detected 96 sequence types (STs) with more than 170 distinct AMR combinations, with isolates from 25 STs harbouring the carbapenem resistance gene *bla*_IMP-4_ ^11^. Plasmids carrying related *bla*_IMP-4_ genes that have been identified in different members of the *Enterobacteriaceae* among clinical isolates from humans, companion animals and wildlife across Australia have also been detected in a gull-associated *E. coli* belonging to ST216 ^11^. Another study using faecal swabs from seagulls in Western Australia detected high frequencies of CIA-R *E. coli* at various geographic locations, ranging from 34.3% to 84.3%, with CIA-R *K. pneumoniae* ranging from 12.5% to 50.0% ^12^. Although the latter study covered only a small proportion of the Australian seagull habitat, there was a strong association between anthropogenic activities and carriage of resistant bacteria by seagulls. These findings support the hypothesis that seagulls largely acquire AMR bacteria from humans, most likely through scavenging activity at sites such as rubbish dumps, sewage plants and intensive farms. Subsequently these bacteria may be transmitted back to humans through faecal contamination of food products or water. The current study aimed to test the hypothesis that AMR transmission is predominantly in the direction from humans to seagulls. In this scenario, if seagulls live at sites away from anthropogenic activity they should have a low carriage of resistant bacteria and less frequent detection of the STs that are commonly found associated to humans. We applied a widely accepted classification system describing geographic remoteness of human habitation in Australia as the measure of likely anthropogenic activity, and highly sensitive and specific laboratory techniques were used to look for associations between human population levels and shedding of CIA-R *E. coli* by seagulls. The phenotypic and molecular characteristics of the target organisms were thoroughly scrutinized to assess whether the resistance types and associated genes were related to those detected in isolates from humans. To maximise the robustness of inferences and their applicability to AMR control programs, this work was conducted on a continental scale, with sampling sites separated by as far as 3,500 km.

## MATERIALS AND METHODS

### Sampling

Seagull habitats around the Australian coastline were sampled between November 2019 and March 2020. Fresh droppings from silver gulls were swabbed from the ground without a preset sample limit, targeting sites from major cities to very remote areas to assess anthropogenic effects on resistant *E. coli.* In total, 81 habitats across 54 localities (up to 3,500 km apart) in Queensland, New South Wales, Victoria, South Australia, and Western Australia were sampled. Sites were assigned to Australian Bureau of Statistics (ABS) ^13^ remoteness categories using Accessibility and Remoteness Index of Australia (ARIA+) ^14^, with mean ARIA+ used as a quantitative remoteness measure. The five remoteness categories (MCA, IRA, ORA, RA and VRA, with increasing remoteness) are defined in **Table 1** together with sampling numbers from these sites. Swabs were collected into Ames charcoal media (Copan, USA), transported on ice, and processed within five days at the laboratory.

**Table 1.** Sampling from seagull habitats: Australian Bureau of Statistics remoteness categories, range and mean of ARIA+ values quantifying degree of remoteness, number of geographic sites from which seagull faecal samples were obtained, and number of samples obtained.

| Remoteness category | Name | ARIA+ value ranges (min - max) | Fixed ARIA+ value | No. sites / No. samples |
| --- | --- | --- | --- | --- |
| 0 | Major cities of Australia (MCA) | 0 / 0.20 | 0.10 | 21 / 388 |
| 1 | Inner regional Australia (IRA) | 0.21 / 2.40 | 1.30 | 26 / 471 |
| 2 | Outer regional Australia (ORA) | 2.39 / 5.92 | 4.16 | 10 / 174 |
| 3 | Remote Australia (RA) | 5.93 / 10.53 | 8.23 | 14 / 223 |
| 4 | Very remote Australia (VRA) | 10.54 / 15.00 | 12.77 | 10 / 160 |

**ARIA+ is an official Australian Bureau of Statistics rating (on a scale of 0 to 15) of the geographic remoteness of locations based on the distance from service centres.**

### Bacterial isolation and species identification

The swabs were enriched by placing the tips in three mL of buffered peptone water for four hours at 37°C. The enriched samples were streaked onto three different selective agar plates designed to isolate *E. coli* with resistance to CIAs. These included Brilliance ESBL agar, MacConkey agar infused with 1 mg/L ciprofloxacin and Brilliance CRE agar. The plates were incubated for 16 – 40 hours at 37°C. Presumptive *E. coli* isolates (one colony per plate) were sub-cultured on Sheep Blood agar plates and identified by MALDI-TOF MS. Resistant isolates identified as *E. coli* were used for further study ^10^.

### Antimicrobial susceptibility testing

Minimal inhibitory concentrations for the CIA-R isolates were determined using an antimicrobial panel consisting of 13 antimicrobials that covered 8 antimicrobial classes. These included amikacin, ampicillin, cefotaxime, ceftazidime, chloramphenicol, ciprofloxacin, colistin, florfenicol, gentamicin, meropenem, sulfamethoxazole, tetracycline, and trimethoprim. CIAs included amk, amp, cta, ctz, cip, col, gen and mer. Broth microdilution was performed using the Robotic Antimicrobial Susceptibility Platform (RASP) ^15^, based on Clinical Laboratory Standards Institute (CLSI) 30^th^ edition M100 guidelines ^16^, and the breakpoints were interpreted using EUCAST epidemiological cut-off value (ECOFF). *E. coli* ATCC 25922 was used as the control strain.

### Whole genome sequencing

Whole genome sequencing (WGS) was performed on 218 of the CIA-R *E. coli* isolates. They were selected for WGS if they showed multiclass resistance (MCR) in the MIC assays, had unique phenotypic resistance profiles, and were from different sampling locations to avoid selecting the same clone. Bacterial DNA was extracted from overnight cultures of single isolates on Sheep Blood Agar using the MagMax DNA Multi-Sample extraction kit (Thermo Fisher) according to manufacturer’s protocol. Sequencing libraries were prepared using the Celero DNA-seq library preparation kit (Tecan, Switzerland) according to the manufacturer’s protocol. Sequencing was performed on the Illumina NextSeq 550 platform using a Mid-Output 300 cycles Kit v2.5 (Illumina, US). A subset of isolates underwent Oxford Nanopore sequencing on the PromethION platform to determine location of the resistance determinants.

### Genomic Analysis and Characterization

Raw reads were processed using the Nullarbor pipeline ^17^, and subjected to *de novo* assembly via SPAdes v3.14.0 ^18^, checking the quality using QUAST v5.3.0. Multilocus sequence type (ST in MLST) for each isolate was identified using the mlst tool (v2.19.0) based on the pubMLST database ^19^. Virulence genes and serotypes were identified in assembled draft genomes using ABRicate v0.8 with the databases VFDB ^20^ and EcOH, respectively, based on a 90% coverage and identity cut-off. *E. coli* and FimH typing (ST131) were carried out using FimTyper v1.0 ^21^. Plasmids were identified using ABRicate with the PlasmidFinder database ^22^ with 90% identity and coverage. Acquired and point-mutation antimicrobial resistance (AMR) genes were identified using the abriTAMR pipeline v1.0.19 and the standard parameters ^23^. The genomic data have been deposited in the European Nucleotide Archive under accession number PRJEB121166.

### Major ST phylogeny and SNP analysis

STs determined from this collection were tabulated in Stata v18.0 (StataCorp LLC, TX, USA). The five most predominant STs that all included 10 or more isolates, were identified and related available genomic data for each were downloaded from EnteroBase (https://enterobase.warwick.ac.uk/) for phylogenetic comparisons. Genomic data from EnteroBase included isolates from farm animals, wild animals, companion animals, humans and environmental samples, and only if information of host, geographical origin and year of sampling was available. The data were analysed for core genome single nucleotide polymorphisms (SNPs) using the NASP v1.1.0 pipeline, with the following genomic references for each of the main circulating STs: (NCBI accession numbers) for ST10 (CP031214.1), ST38 (NZ_CP023364.1), ST131 (NZ_CP103672.1), ST744 (JAQQAX000000000.1), and ST1193 (NZ_CP058900.1) and removing recombinations using Gubbins ^24^. Phylogenetic trees were constructed using ModelFinder and 100 bootstrap replicates with IQ-TREE v2.1.2 ^25^, and annotated using Interactive Tree of Life (iTOL) v6 ^26^ Genotypic data (ST) were combined with phenotypic resistance data, and the ten most common STs were selected for downstream visualisation. These STs were selected based on the highest overall prevalence, and, if STs had the same prevalence, then the STs present in more Australian states were selected.

### Statistical analysis

The isolates recovered on the three different selective isolation plates were divided into four categories: resistant to ESBL, resistant to CIP, resistant to CRE, and resistant to any CIA. An assessment of association between remoteness of sampling and the proportion of seagulls found shedding these categories of CIA-R *E. coli* was performed in Stata v18.0. Data were fitted to a mixed effect generalized linear model (meglm command in Stata) with binomial error distribution and logit link function. Fixed effects comprised the mean ARIA value for the remoteness category and resistance phenotype (comprising resistance to CIP, CRE, ESBL and CIA), and with random effects being resistance phenotype within location. From the fitted model, marginal predictions and 95% confidence limits were generated for each combination of resistance phenotype and for each fixed ARIA+ value corresponding to the mean ARIA+ for each remoteness category. Inference was based on estimation rather than significance testing, with the magnitude and precision of these effect estimates, expressed as 95% confidence intervals, used to assess the strength of evidence for an effect of remoteness on resistance prevalence.

## RESULTS

### CIA-R *E. coli* and their geographic occurrence

Of the 1,416 faecal samples collected, presumptive CIA-R *E. coli* growth on the selective agar was observed for 567 (39.8%). The numbers of samples with growth for each selective agar were 271 (19.0%) from Brilliance ESBL plates, 212 (14.8%) from MacConkey + ciprofloxacin agar plates, and 84 (5.9%) from Brilliance CRE plates. **Figure 1** presents the percentage of resistance phenotypes from various classified area categories together with the corresponding mean. The percent of *E. coli* resistant to CIP were generally lower in more remote areas, with the highest percentage (21.2%) being in IRA and the lowest in ORA (3.4%). ESBL-producing *E. coli* were less in ORA (15.7%) and VRA (0.8%). For CRE-resistant *E. coli*, there was a slight trend towards less resistance in more remote areas (8.5% in MCA and 0% in VRA). Overall, CIA-R *E. coli* were less in ORA (9.9%) and VRA (4.7%). **Figure 2** represents the effect of remoteness on predicted percentage of *E. coli* belonging to four different resistance phenotypes. For each of the four phenotypes, a trend for decreased prevalence of resistance with increased remoteness was observed, with the accompanying 95% confidence intervals used to assess the certainty of this pattern for each phenotype (**Figure 2**).

**Figure 1.**
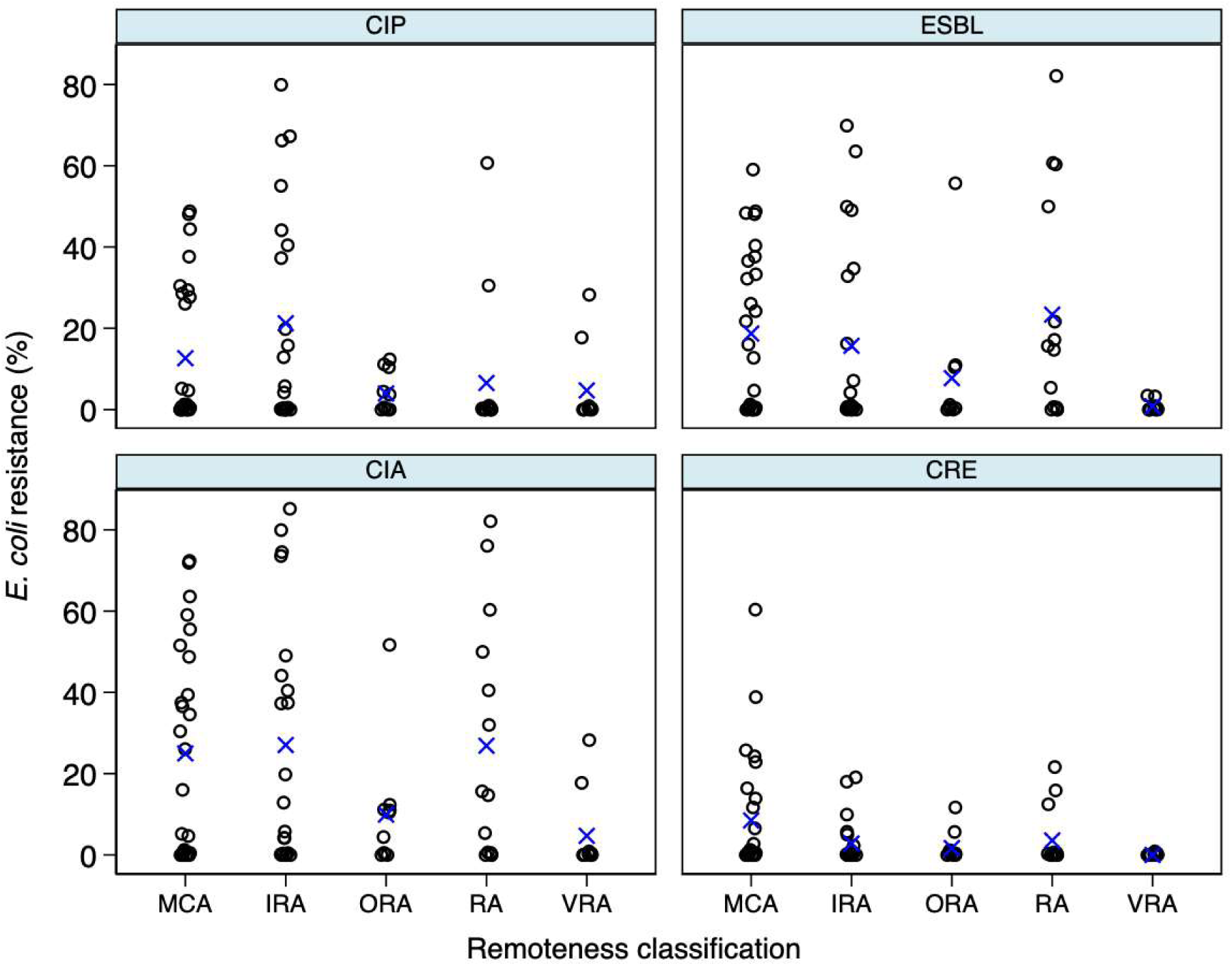
Proportion of *E. coli* isolates exhibiting four antimicrobial resistance phenotypes— ciprofloxacin-resistant (CIP), extended-spectrum β-lactamase–producing (ESBL), carbapenem-resistant (CRE), and resistant to any critically important antimicrobial (CIA)—across Australian Statistical Geography Standard (ASGS) remoteness categories. [Black circles represent individual sampling sites within each category, and blue “×” symbols denote category means. Remoteness categories, in increasing order of remoteness, include: Major Cities of Australia (MCA), Inner Regional Australia (IRA), Outer Regional Australia (ORA), Remote Australia (RA), and Very Remote Australia (VRA)].

**Figure 2.**
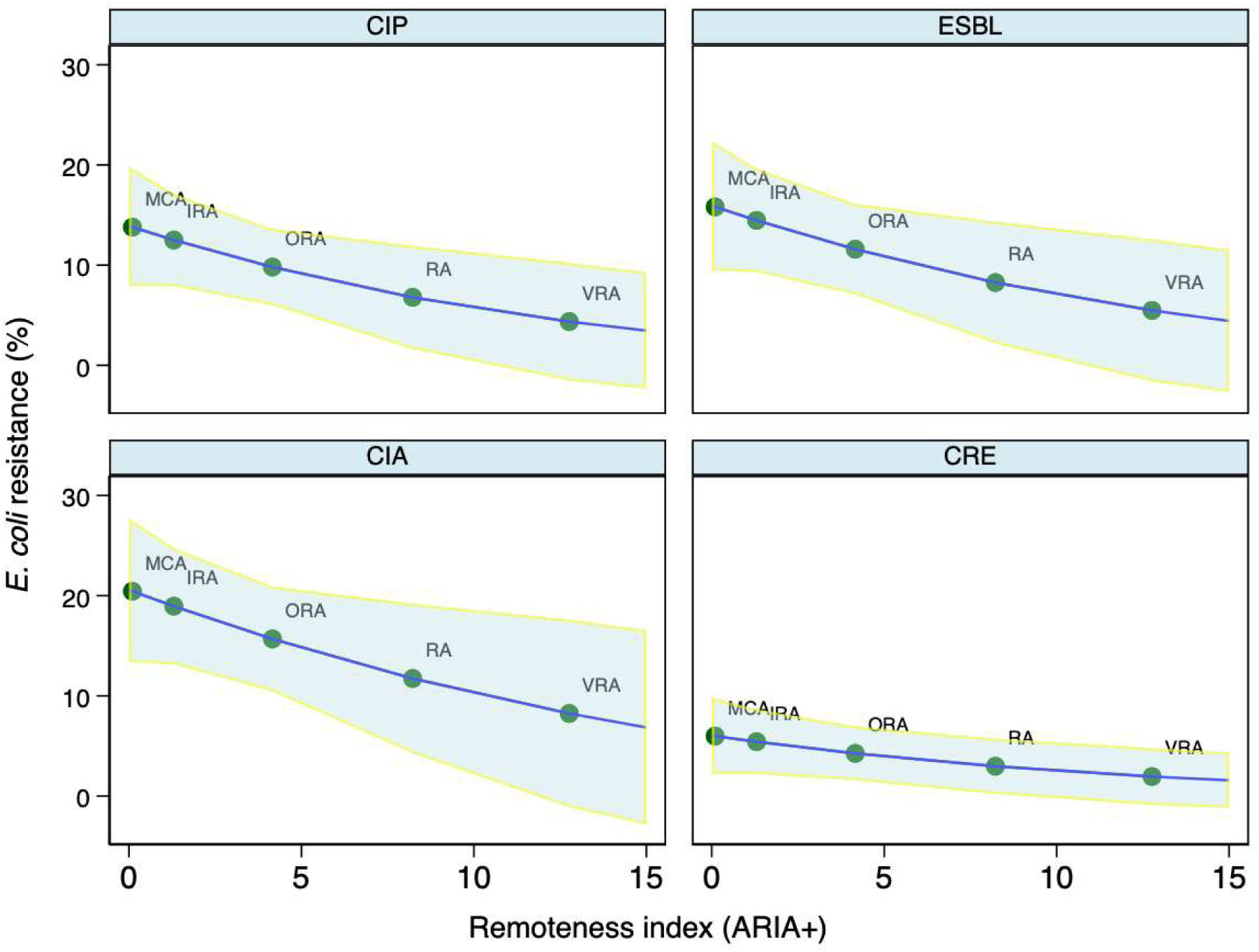
Output from mixed effects generalized linear model showing the effect of remoteness of sampling (ARIA+) on predicted percent of *E. coli* (with 95% confidence intervals) belonging to each of four resistance phenotypes (CIP: Ciprofloxacin; ESBL: Extended-spectrum β-lactamase; CRE: Carbapenem; CIA: Any critically important antimicrobial). The mean ARIA+ value for each category have been annotated. Abbreviations: Major Cities of Australia (MCA), Inner Regional Australia (IRA), Outer Regional Australia (ORA), Remote Australia (RA), and Very Remote Australia (VRA).

### MIC results

Amongst the 567 isolates subjected to MIC testing, a high prevalence of resistance was observed to ampicillin (94.0%), ciprofloxacin (87.8%), cefotaxime (73.7%) and ceftazidime (70.7%). Low levels of resistance were found against amikacin (2.5%) and one isolate showed resistance to colistin (**Figure 3**). Phenotypic resistance to various drug classes observed ranged between fully susceptible to a maximum of eight drug classes (**Supplementary Table 1**), and a total of 519 strains were MDR (≥3 antibiotic classes).

**Figure 3.**
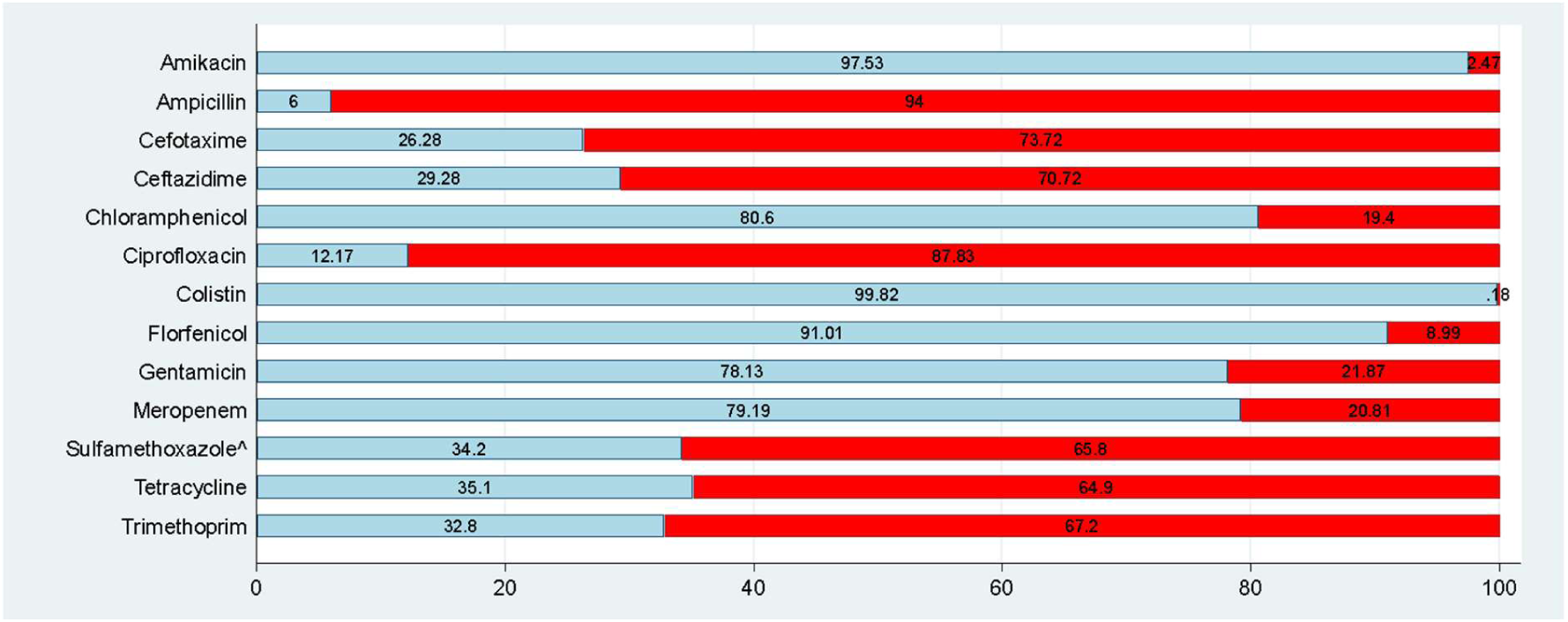
Bar plot displaying from the MIC results. Blue represents percentage of wild type and red represents percent non-wild type, unless otherwise indicated by footnotes. Nil footnote is used for a drug when data can be (and are) interpreted using a wild/non-wild type breakpoint. ^ data represent the percent clinically resistant due to lack of breakpoints for both wild type and susceptible

### Isolates used for WGS, sequence types and resistance genes

From the 567 CIA-R *E. coli* isolates, the 218 selected for WGS consisted of 14 CRE-, 79 CIP-, and 105 ESBL-resistant isolates. Using MLST on the genomic data, these isolates belonged to 61 STs. The most common ST was ST131 (36 isolates, 16.51%). This ST also was the most prevalent across all Australian States, followed by ST1193, ST38, ST744 and ST10 (**Figure 4**). Others were more regional, particularly ST1193, which was only found in South Australia and Western Australia, and ST10 found only in Western Australia. New South Wales had the highest diversity of STs, followed by Victoria. The most prevalent genes encoding resistance detected amongst the 218 isolates included *bla*_TEM_ (53.2%) followed by *bla*_CTX-M-15_ (33.5%), *bla*_CMY-2_ (12.8%), *bla*_CTX-M-27_ (9.2%), and *bla*_CTX-M-14_ (7.8%) (**Supplementary Table 2**). One isolate from Victoria (W’bool Lookout) belonging to ST1011 harboured *mcr1.1* (encoding colistin resistance) and *tet*(X4) (encoding tigecycline resistance). Across 37 AMR gene categories, resistance determinants against β-lactams were detected in most isolates, with 90.0% carrying at least one β-lactam resistance gene or ESBL gene. Further, 40/218 isolates (18.3%) carried AmpC-type β-lactamases (e.g., *bla*_CMY-2_, *bla*_DHA-1_, and *bla*_EC_ variants), while 122/218 isolates (56%) carried extended-spectrum β-lactamases (ESBLs; predominantly *bla*_CTX-M_ family members such as *bla*_CTX-M-15, –27, –55_). Similarly, 96% of isolates showed phenotypic resistance to β-lactams. Further, 117/218 isolates (53.7%) carried genes for narrow-spectrum β-lactamases (e.g., *bla*_TEM_ variants), with 70.7% of isolates showing phenotypic resistance. Quinolone-resistance mutations in the quinolone-resistance-determining regions (QRDRs) of *gyrA* and *parC* were also common. Here, 151/218 (69.3%) of isolates carried at least one *gyrA* substitution (most commonly S83L, D87N), and 137/218 (63%) carried various *parC* and *parE* mutations. These mutations correlated with phenotypic quinolone resistance, with 84.3% of isolates showing phenotypic resistance. Similarly, 149/218 isolates (68%) carried sulfonamide resistance genes (*sul1, sul2, sul3,*and *folP_P64S*), with 97.5% showing phenotypic resistance to folate pathway inhibitors. Trimethoprim-resistance genes were carried by 130/218 isolates (59.6%). For tetracycline, genotypic resistance was detected in 134 isolates (61.5%), and the corresponding phenotypic resistance was 64.6%. Isolates carrying AMR genes to CIAs including colistin, carbapenems, metallo-β-Lactams (NDM), and tigecycline were less common. Colistin-resistance determinants, including *mcr-1*, were identified in a single isolate, which also harboured resistance determinants for quinolones, ESBLs, fosfomycin, aminoglycosides, tetracyclines, tigecycline, sulfonamides, and streptomycin. This isolate also carried the tigecycline resistance gene *tet*(X4). Carbapenemase gene carriage (*bla*_OXA-23,_ ompF_Q88STOP) was rare: only 2 isolates (1.0%) were detected *in silico* and displayed phenotypic resistance, whilst *bla*_NDM-5_ was present in three isolates (1.4%) which also displayed carbapenem resistance. Five isolates (2.3%) carried the *rmt*B gene responsible for high-level aminoglycoside resistance mediated by 16S rRNA methyltransferase. These isolates were also MDR, including to fluoroquinolones and AmpC beta-lactamase (*bla*_CMY-42_). The analysis of AMR gene carriage for the top 10 most common STs and their multiclass resistance phenotype profiles showed considerable variability in resistance traits (**Figure 5**). Here, ST38, ST457, and ST131 exhibited the highest median values for both AMR gene carriage and multiclass resistance, indicating a broader spectrum of resistance mechanisms. The variation in AMR gene carriage for these STs, as indicated by their interquartile ranges, suggested a higher diversity in resistance profiles compared to other STs. In contrast, ST648 and ST963 displayed lower median values and narrower interquartile ranges, indicating more restricted resistance profiles with less variability. Isolates of STs with a greater number of AMR genes tended also to have a higher number of MCR, suggesting a correlation between the number of resistance genes and phenotypic resistance diversity.

**Figure 4.**
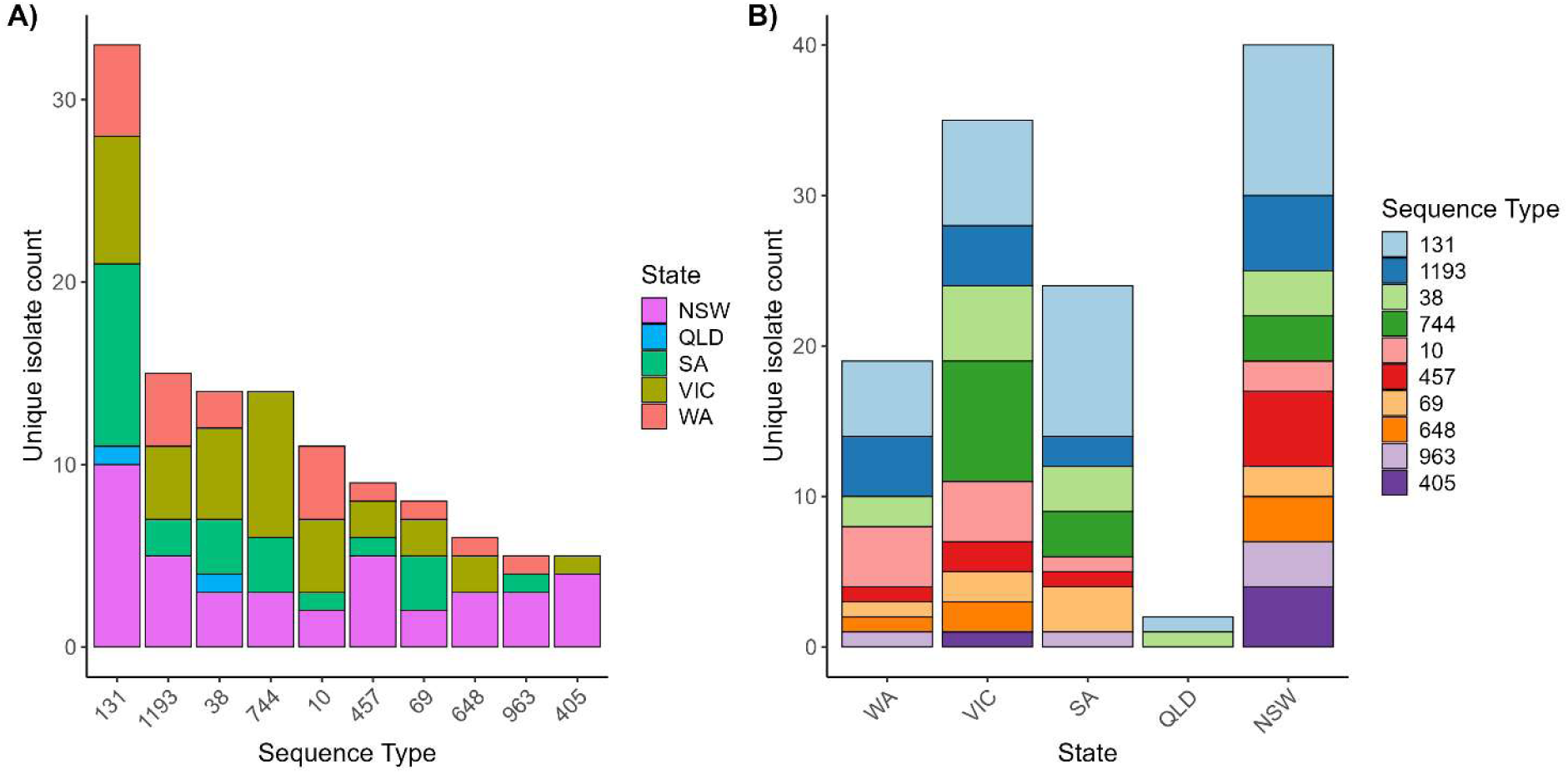
Top ten *E. coli* sequence types (STs) identified among critically important antimicrobial resistant *E. coli* isolated from seagulls across Australian states. A) distribution based on dominant ST, B) distribution based on state of isolation. Abbreviations: Western Australia (WA), Victoria (VIC), South Australia (SA), Queensland (QLD), and New South Wales (NSW)

**Figure 5.**
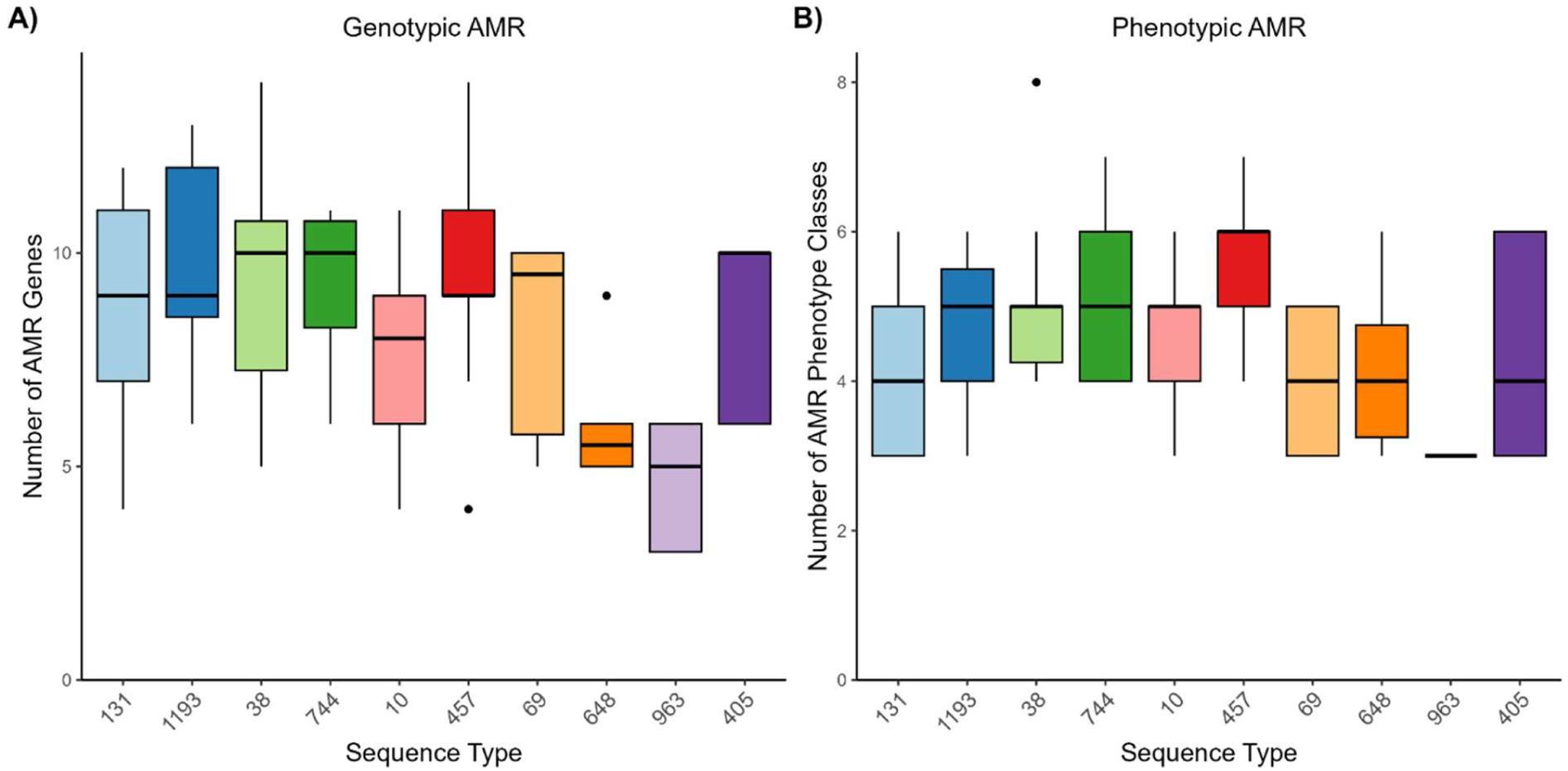
Phenotypic and genotypic AMR of the 10 most prevalent STs identified among CIA-R *E. coli* isolates from seagulls across Australia. A) distribution based on genotypic AMR, B) distribution based on phenotypic AMR

### Virulence genes

Virulence gene characterization revealed that the most frequently detected STs carried virulence genes associated with extra intestinal infections (**Supplementary Table 3**). Among the ten most common STs, ST38, ST457 and ST131 exhibited the highest median counts and widest inter-quartile ranges of virulence genes, whereas ST744, ST963 and ST10 showed both lower medians and tighter ranges (**Figure 6**). At the gene level, 45.5% of isolates carried the serum-survival factor *iss* and 61.5% harbored the yersiniabactin synthesis genes *ybtP/Q*. Genes encoding the aerobactin (*iutA*, 45.9%) and salmochelin (*iroN*, 7.8%) receptors also were present. The classical UPEC adhesins such as *papGII* (8.7%) and *sfaF* (0.5%) were rare, together with toxins *hlyA* (4.6%) and *cnf1* (2.3%). The autotransporter *fdeC* occurred in 95.4% of isolates.

**Figure 6.**
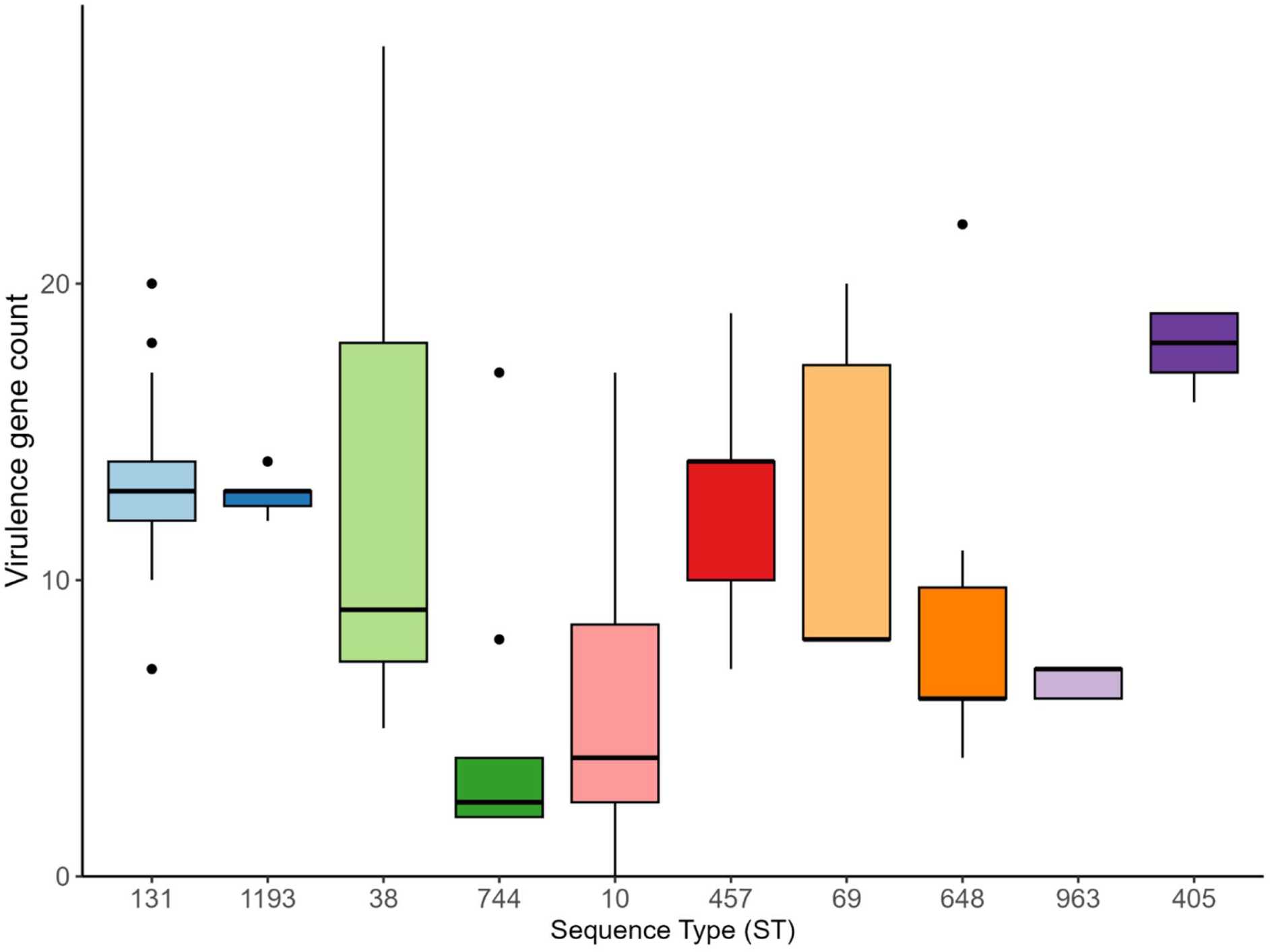
Difference in virulence genes count carried by the top 10 most common sequence types identified among the seagulls sampled across Australia.

Pandemic extraintestinal pathogenic *E. coli* (ExPEC) lineages ST131 and ST1193 uniformly possessed *iss*, *fdeC*, *ybtP/Q*, *iutA*, *iroN*, and *fimH*. ST744 likewise retained this full ExPEC gene toolkit, supporting robust serum resistance, iron scavenging and adhesion. By contrast, ST648 showed scant carriage of *iss* and yersiniabactin genes, suggesting a lower virulence potential, while ST38 presented an intermediate profile with consistent *fimH*, *iss* and *ybtP/Q* plus frequent *iutA/iroN*. ST963 contained the sparsest virulence set, lacking most key ExPEC markers and thus resembling a commensal-like genotype.

### Plasmid, serotype and *fim*H distribution

A wide variety of plasmids and serotypes were identified in the sequenced collection, with several isolates carrying multiple plasmids. IncFIB were the most abundant (60.6%), followed by Col156 (40.4%) mostly associated with ST131, 1193 and 648, and IncFIA (38.1%) associated with STs 131, 1193 and 405 (**Supplementary Table 4**). Several isolates did not have their O or H antigens identifiable *in silico*, but those identified had more commonly occurring serotypes Onovel31:H4 (N=23 occurrences), O75:H5 (N=14), and O16:H5 and O11:H25 (N=13 and 11 respectively) (**Supplementary Table 5**). FimH was tested only on ST131 isolates given the usual connection of certain *fim*H subtypes to particular hosts, with the majority of the ST131 isolates belonging to the H30 subtype (N=16/36) followed by H41 (N=13/36) (**Supplementary Table 6**).

### Phylogenetic analyses

Despite most EnteroBase genomes for the selected STs (ST10, 38, 131, 744, and 1193) belonging to human isolates, there was a tendency for other sources intermingling in human-dominated clades, with the exception of ST744 where the majority of isolates belonged to wild animals, with a few human isolates clustering among these (**Figure 7**). For each of the main STs, except for ST10, fluoroquinolone resistance (either point mutation or acquired) was overwhelmingly present among all isolates from all sources, particularly in ST1193. ESBL resistance was widespread among all STs and sources except for ST744 of animal origin (**Figure 7-11**). Carbapenem resistance was predominant among ST38 where it was strongly linked to human isolates, and rarely circulating in the other STs, though present in one ST744 and one ST10 isolates from seagulls from this study (**Figure 7-8)**.The pairwise core-genome SNP comparison among each of the main STs and the EnteroBase isolate collections showed high similarities among some isolates from seagulls and humans from Australia in ST10 (5 SNPs), ST131 (<15 SNPs), and ST1193 (15 SNPs).

**Figure 7.**
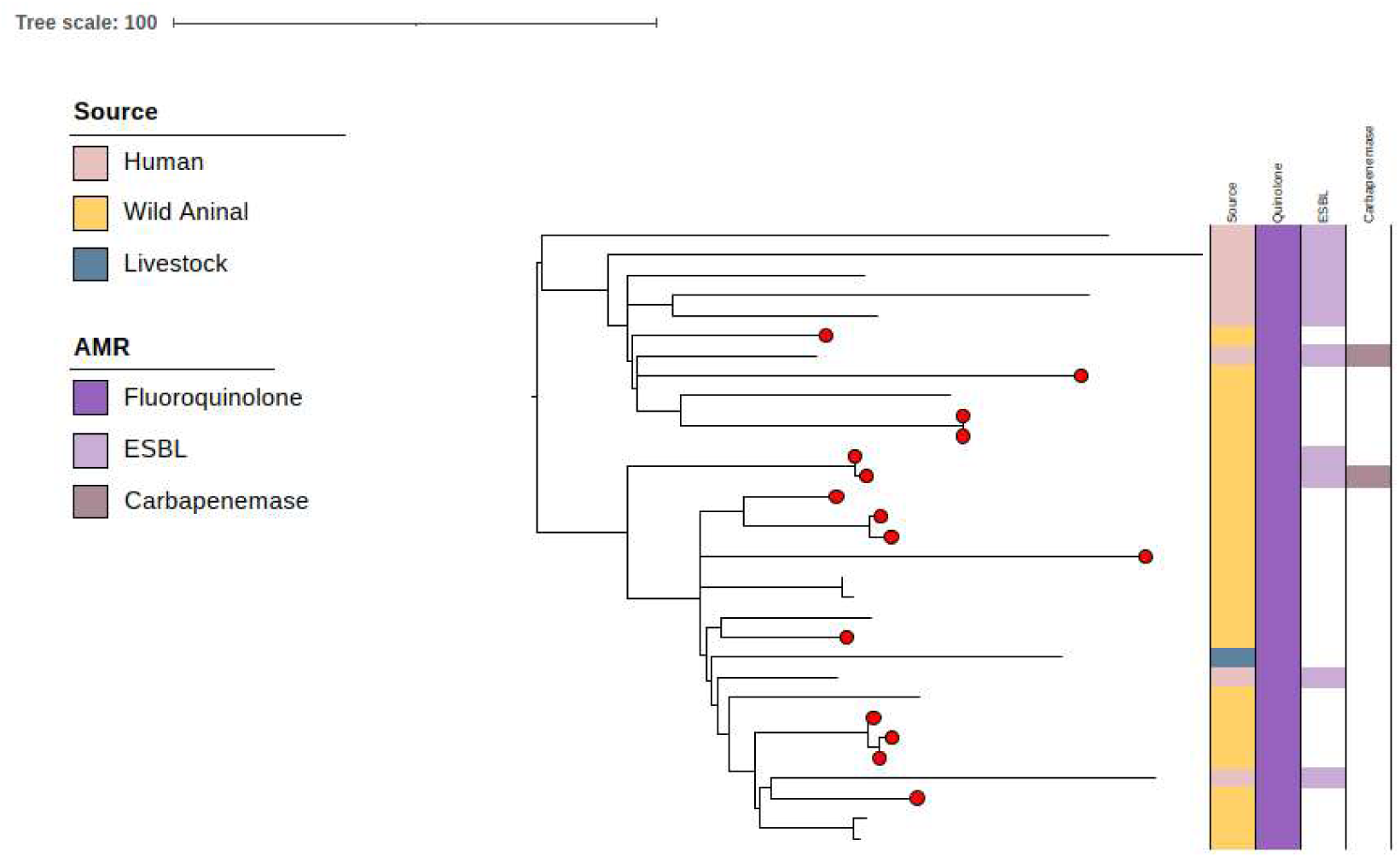
ST744: Core genome SNP phylogeny of ST744 isolates (N=31) and available metadata for source and selected antimicrobial resistance (AMR). Mid-point rooted phylogeny constructed on SNP calling in 79% (4.1Mb) of the reference chromosome. The seagull isolates from this study are highlighted with a red circle at the tip of each branch.

**Figure 8.**
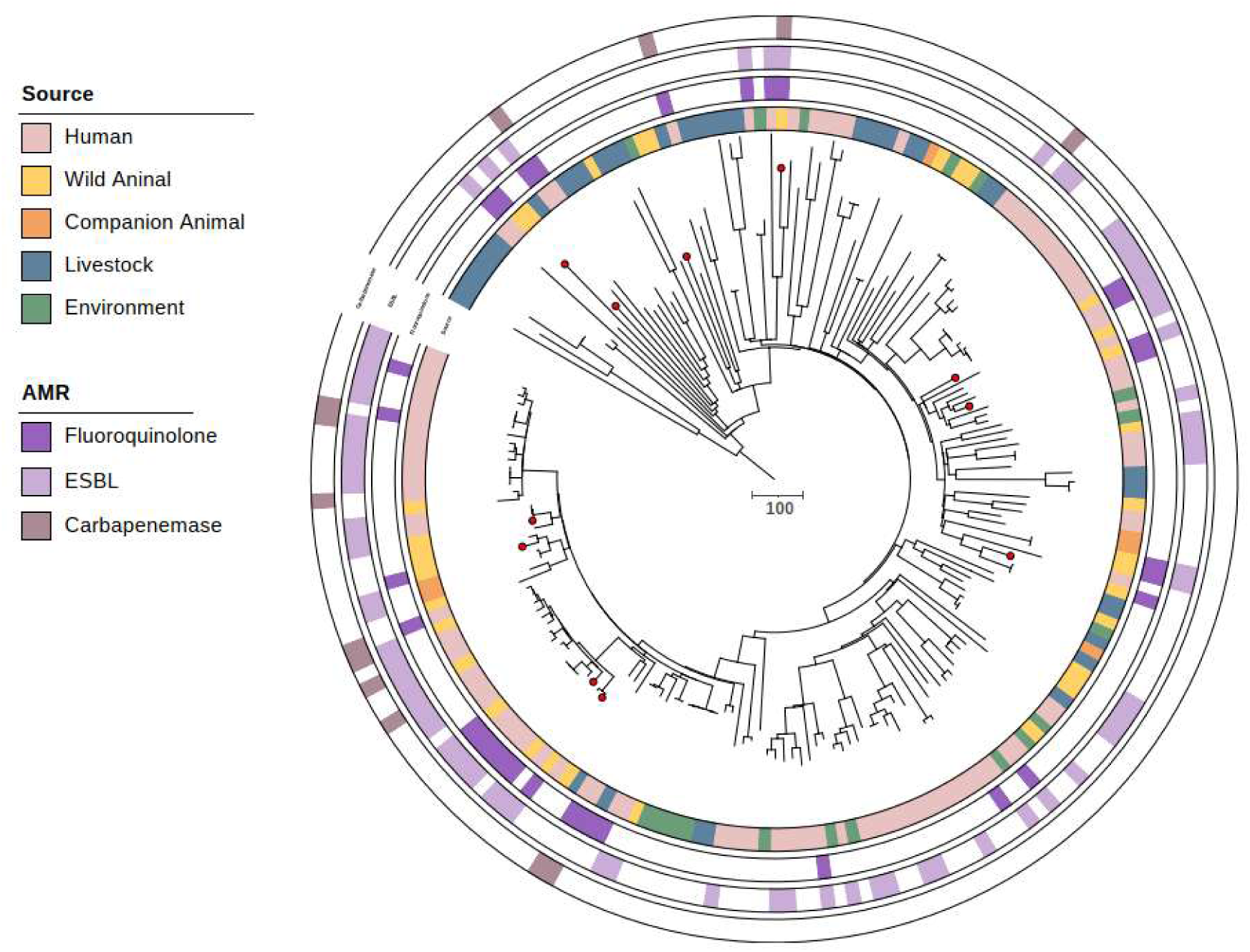
ST10: Core genome SNP phylogeny of ST10 isolates (N=201) and available metadata for source and selected antimicrobial resistance (AMR). Mid-point rooted phylogeny constructed on SNP calling in 64% (2,9Mb) of the reference chromosome. The seagull isolates from this study are highlighted with a red circle at the tip of each branch.

**Figure 9.**
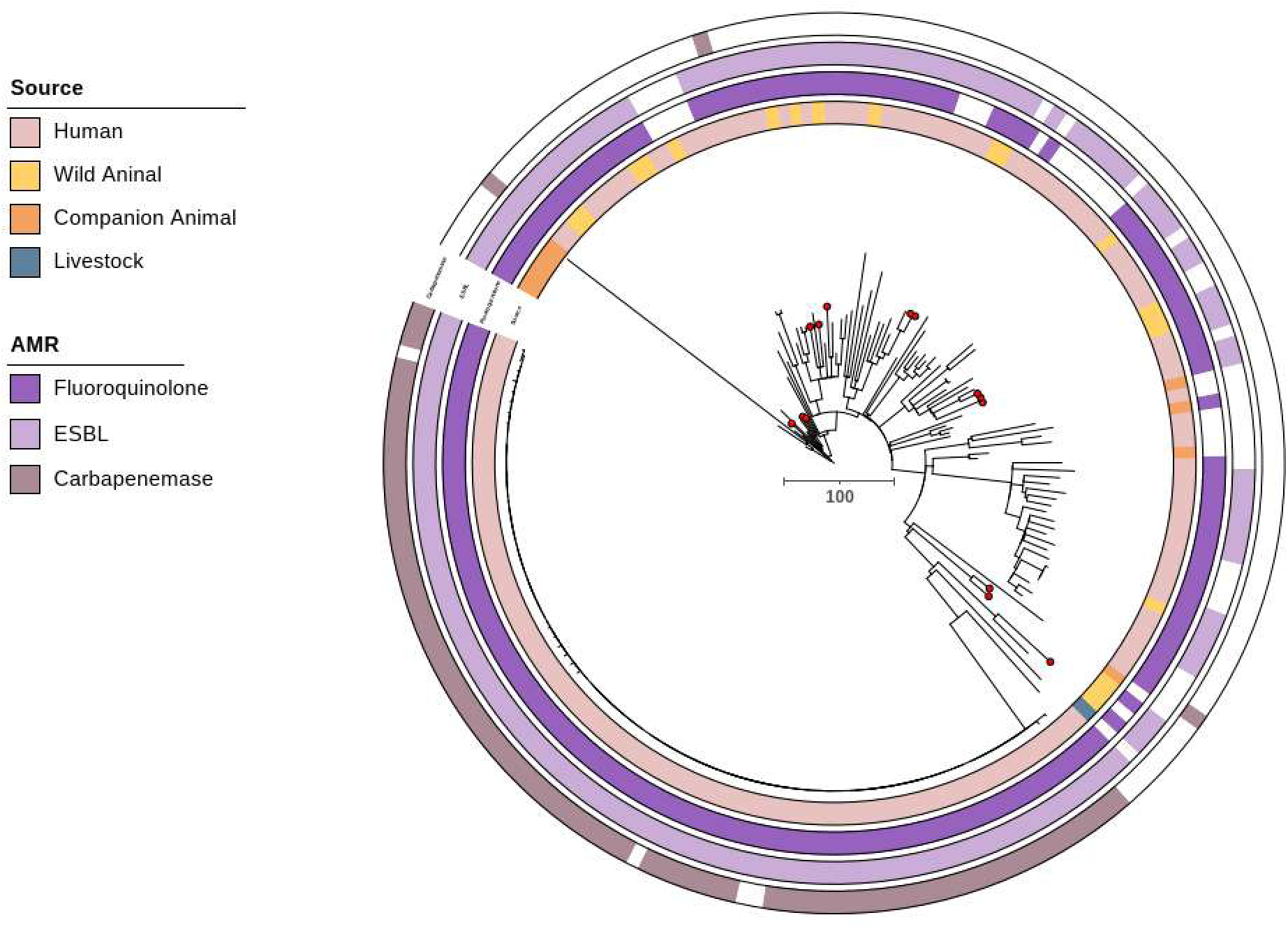
ST38: Core genome SNP phylogeny of ST38 isolates (N=190) and available metadata for source and selected antimicrobial resistance (AMR). Mid-point rooted phylogeny constructed on SNP calling in 75% (3,9Mb) of the reference chromosome. The seagull isolates from this study are highlighted with a red circle at the tip of each branch.

**Figure 10.**
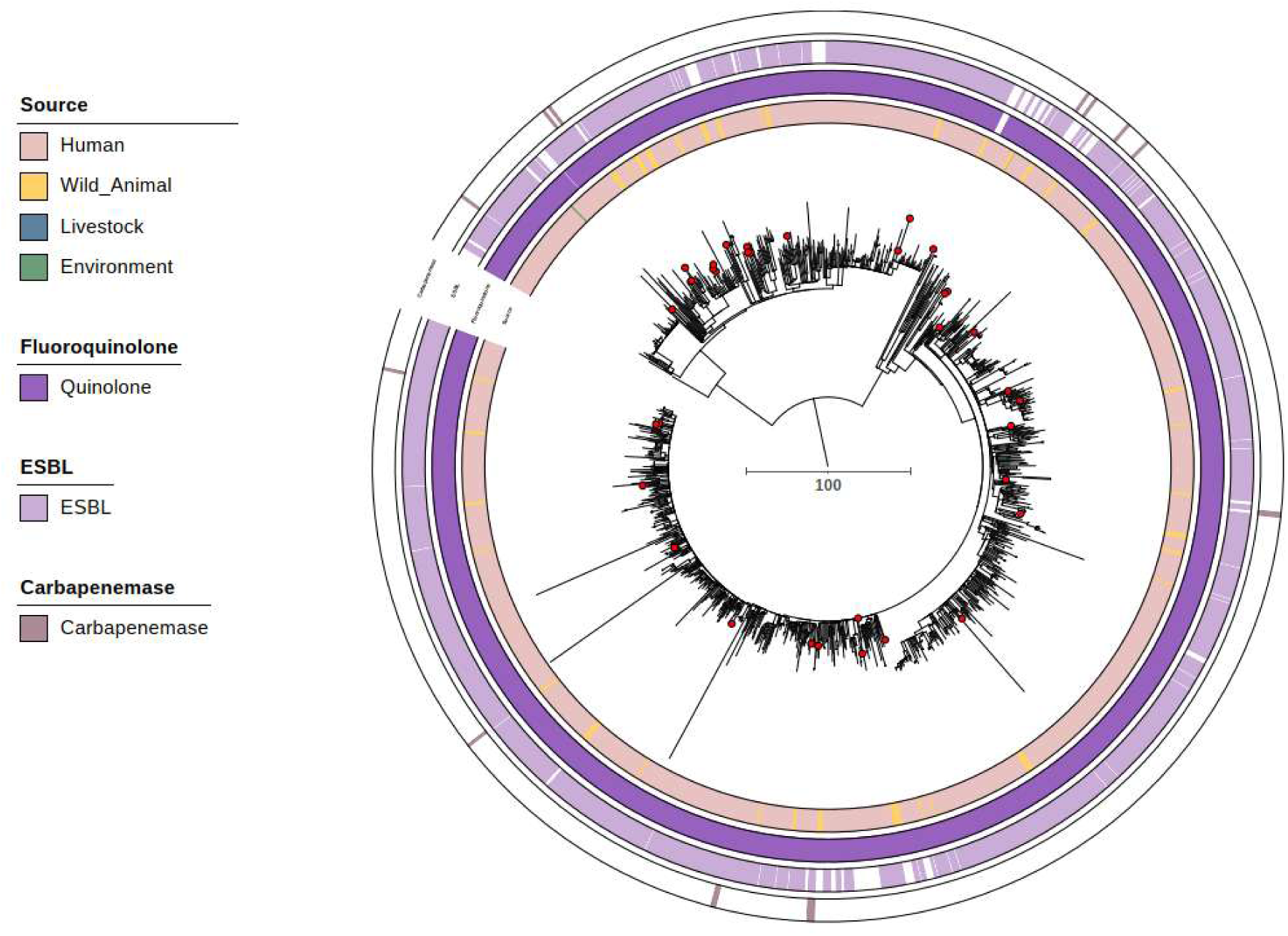
ST131: Core genome SNP phylogeny of ST131 isolates (N=1,199) and available metadata for source and selected antimicrobial resistance (AMR). Mid-point rooted phylogeny constructed on SNP calling in 51.7% (2,8Mb) of the reference chromosome. The seagull isolates from this study are highlighted with a red circle at the tip of each branch.

**Figure 11.**
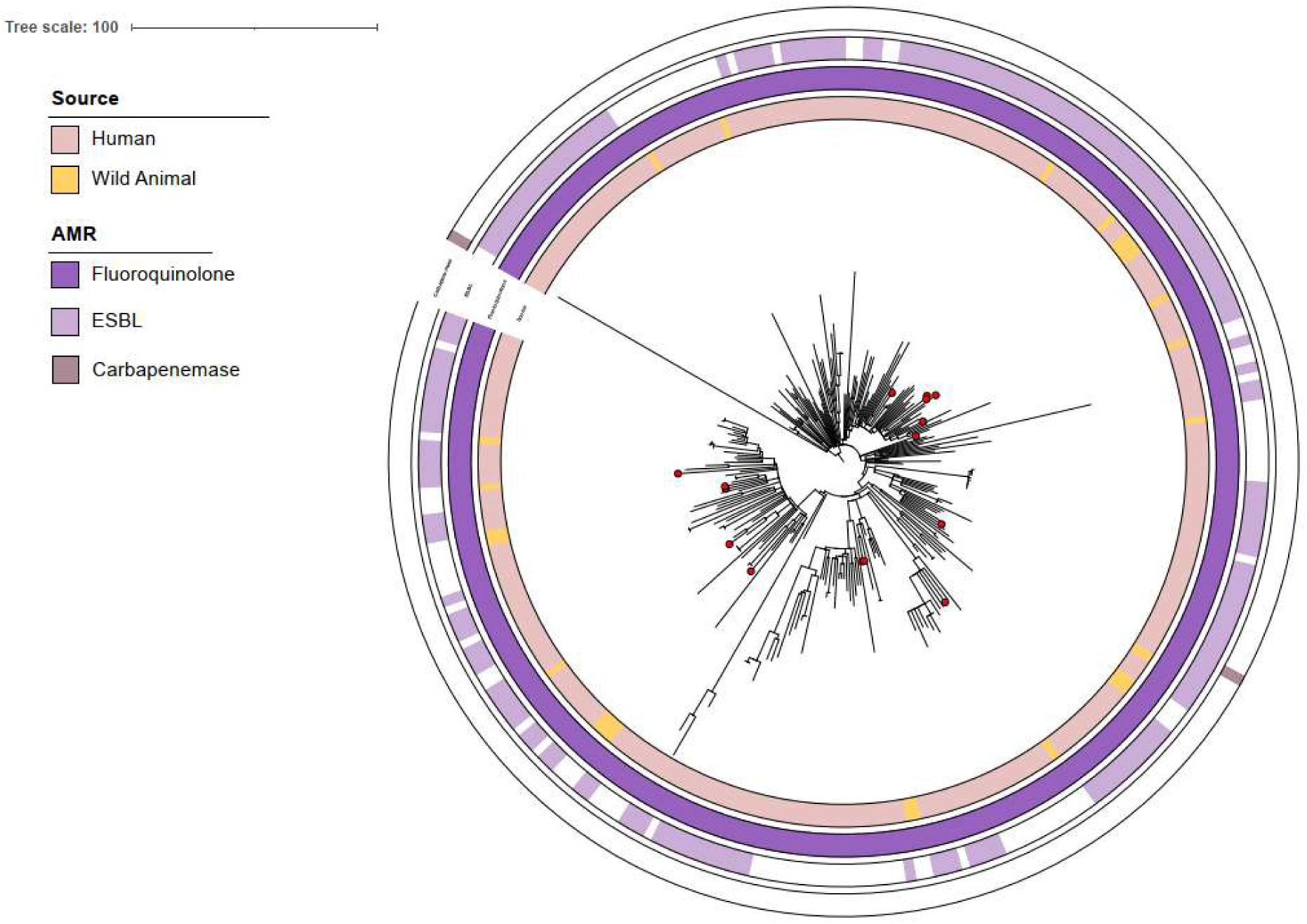
ST1193: Core genome SNP phylogeny of ST1193 isolates (N=277) and available metadata for source and selected antimicrobial resistance (AMR). Mid-point rooted phylogeny constructed on SNP calling in 83% (4.2Mb) of the reference chromosome. The seagull isolates from this study are highlighted with a red circle at the tip of each branch.

## DISCUSSION

This study aimed to investigate the relationship between carriage of CIA-resistant *E. coli* by seagulls and their habitat in relation to human population density. It provides a large-scale genomic and phenotypic analysis of *E. coli* isolated from Australian silver gulls using a high-resolution approach that parallels contemporary livestock AMR surveillance. By sampling gulls across regions, we showed AMR was most common in areas with the greatest human population density and activity. Resistance displayed a clear geographic gradient, decreasing with increasing remoteness. CIA-R *E. coli* were least frequent in ORA (9.9%) and VRA (4.7%), while CIP resistance was highest in IRA (21.2%) and lowest in ORA (3.4%). ESBL-producing *E. coli* showed a similar distribution, with the lowest prevalence in ORA (15.7%) and VRA (0.8%) (**Figure 1**). Consistent with these observations, model-based estimates confirmed a progressive decline in all four resistance phenotypes with increasing distance from human populations (**Figure 2**). This is likely to be due to greater anthropogenic antibiotic selection and exposure in densely populated areas compared with remote, less populated regions, and driven by exposure to municipal wastewater and hospital effluents, or by food-waste facilities and landfills that attract silver gulls ^10, 27–29^. CRE resistance is likely primarily to be hospital-associated and thus shows a less prominent pattern, whereas the prevalence of CIP resistance and ESBL phenotypes may be driven more by other exposure routes which become less common in more remote areas.

The high prevalence of CIA resistance in gull-derived *E. coli* was striking in the context of Australia’s livestock AMR status. National surveillance of Australian meat chickens and pigs consistently shows very low levels of resistance to fluoroquinolones, third-generation cephalosporins and colistin ^31–31^. In contrast, gull isolates in our study frequently carried resistance to both classes, with many exhibiting co-resistance to multiple CIAs. This disparity shows that while Australian livestock production benefits from stringent antimicrobial restrictions, wildlife that interface closely with human environments may serve as uncontrolled reservoirs of high-risk AMR. From a One Health perspective, this finding is concerning because it represents a potential bypass of the very biosecurity and stewardship measures that have kept AMR levels low in the food production sector. Notably, because these CIAs are not used in Australian livestock production, the establishment and proliferation of such CIA-resistant clones in that setting is for now likely to be limited. Genomic characterisation of *E. coli* isolated from gulls further supports the anthropogenic origin of their resistance profiles. The most frequent STs identified in gull isolates, including ST131, ST354 and ST10, are globally distributed human-associated lineages linked to ExPEC infections ^32–33^. Many carried plasmids belonging to incompatibility groups IncF and IncI, which are dominant vehicles for *bla*_CTX-M_ ESBL genes and plasmid-mediated quinolone resistance genes in human clinical isolates ^34–36^. Comparative analysis with Australian livestock datasets shows a clear divergence: these high-risk plasmids and human-associated STs are rare in *E. coli* from livestock, where resistance is largely confined to tetracycline and ampicillin determinants ^37–38^. The close genomic similarity between some STs belonging to gull isolates and human clinical strains such as in the case of ST10, ST131 and ST1193, combined with the gulls’ reliance on anthropogenic food sources such as landfill and wastewater outfalls ^10, 39, 40^, strongly suggests that human waste streams are a primary source of these AMR clones. This was also evidenced in ST131 isolates, where *fim*H subtypes H30 and H41, which are strongly associated with human clinical isolates ^41^, were predominant in *E. coli* from seagulls. The detection of multiple resistance determinants to CIAs in globally disseminated lineages has direct public health implications. ST131, a pandemic lineage and dominant cause of multidrug-resistant urinary tract and bloodstream infections in humans, was prominent among the gull isolates, often carrying *bla*_CTX-M-15_ or *bla*_CTX-M-27_ ^42^. The co-occurrence of *qnrS* and *aac(6′)-Ib-cr* with chromosomal quinolone resistance mutations mirrors patterns seen in hospital-derived isolates ^43, 44^, indicating that gulls may be acquiring fully equipped MDR strains rather than assembling them *in situ*. The ability of these clones to persist in the gut microbiome of gulls in the absence of antimicrobial selection pressure, as shown for ST354 in Australian poultry, could mean that they are maintained and disseminated over long distances. Australian seagull species can travel 10–30 or more km daily and can disperse over long-ranges (80-1,000 km) ^45, 46^, potentially enabling inter-regional or international spread of resistant clones acquired from spending prolonged periods near human populations ^47–49^. Alternatively, seagulls carrying resistant clones from distant human populations could introduce those clones into local Australian human populations. From an ecological standpoint, gulls may function as AMR dissemination hubs. Their opportunistic feeding habits expose them to a diverse array of bacterial communities present in human waste, livestock manure, and natural environments. Once acquired, resistant strains can be redistributed through fecal deposition at roosts, waterways, agricultural fields and coastal environments, facilitating cross-ecosystem transfer ^50, 51^. Studies from Europe, North America and Asia all consistently report a high prevalence of CIA-resistant *E. coli* and *Klebsiella pneumoniae* in gulls, with genetic profiles indistinguishable from human clinical isolates ^10, 49^. Our findings align with this global trend, but highlight a critical surveillance gap: wildlife, and particularly urban-foraging birds, are not included in Australia’s national AMR monitoring frameworks. Given their demonstrated capacity to connect human, animal and environmental reservoirs, their omission represents a blind spot in current risk assessment models. A limitation of our study is its cross-sectional design, which captures a single timepoint and may not reflect seasonal or inter-annual variation in AMR prevalence. Sampling was also restricted to silver gulls, and while they are the dominant urban gull species in Australia, the AMR dynamics in other bird species and non-urban gull populations may differ. Furthermore, while genomic comparisons indicate strong similarity to human-derived isolates, our study did not perform source attribution modelling or include direct environmental sampling from gull foraging sites, which would strengthen causal inference. Despite these limitations, the large sample size, high-resolution genomic analysis and integration of comparative livestock and human AMR datasets provide a robust foundation for interpreting the public health significance of our findings. In conclusion, this study demonstrates that the *E. coli* that Australian silver gulls carry in their intestinal tracts have high rates of resistance to CIAs, in stark contrast to the low levels observed in Australian livestock. The frequency of carriage of resistant *E. coli* by the gulls increased with human population density, consistent with transmission primarily from humans to gulls. The resistance gene content and ST distribution in gull isolates closely mirrored those in human clinical strains, indicating they are likely to have been acquired by gulls from anthropogenic sources and highlighting the role of gulls as mobile reservoirs and potential vectors of high-risk AMR. The ecology and mobility of gulls give them the capacity to bridge otherwise biosecure compartments, potentially undermining national AMR containment strategies. Inclusion of urban wildlife, particularly gulls, in integrated AMR surveillance programs would close a critical gap and provide early warning of emerging resistance threats moving through the environment. Addressing this blind spot will be essential to maintaining Australia’s favourable livestock AMR status and protecting public health in an increasingly interconnected One Health landscape.

## SUPPLEMENTAL MATERIAL

**Supplementary Table 1.** Phenotype profiles and the number and percent of isolates belonging to each phenotype.

**Supplementary Table 2** Genotypic antimicrobial resistance profiles of CIA-Resistant *E.coli* Isolates recovered from Australian seagulls.

**Supplementary Table 3** Virulence gene repertoires of *E. coli* isolates recovered from Australian seagulls.

**Supplementary Table 4** Plasmid replicon types and sequence type associations identified in Australian seagull-derived *E. coli* genomes.

**Supplementary Table 5** Serotype distribution of CIA-resistant *E. coli* isolates from Australian seagulls.

**Supplementary Table 6** FimH allelic subtype profiles of ST131 *E. coli* isolates from seagulls.

## DATA AVAILABILITY

The genomic data have been deposited in the European Nucleotide Archive under accession number PRJEB121166.

## ACKNOWLEDGEMENTS

The authors would like to acknowledge Gaynor Schultz (University of the Sunshine Coast) for assistance with sample collection. We also thank Dr. Susan Lean, John Blinco, Dr. Terence Lee, Dr. Soraya Leedham, and Dr. Ali Harb from the Antimicrobial Resistance and Infectious Disease Laboratory, Murdoch University, for technical assistance with various aspects of data capture for this manuscript.

## Transparency declarations

Financial conflicts of interest and the funder (s) – None to declare for all authors. AI disclosure: We used ChatGPT (OpenAI) solely for grammar, spelling, and stylistic clarity. The authors wrote the manuscript and performed all intellectual editing; no content was generated by the tool, and the authors are responsible for all text.

## Funding

This research was supported by WA Near Miss Fellowship through the Future Health Research and Innovation Fund (FHRI) (Grant No. WANMA/EL2022/10).

